# No evidence for avoidance of black rat scent by the presumably less competitive Natal multimammate mouse in a choice experiment

**DOI:** 10.1101/103853

**Authors:** Laura N. Cuypers, Wim L. Cuypers, Amélie Gildemyn-Blomme, Laura Abraham, Senne Aertbeliën, Apia W. Massawe, Benny Borremans, Sophie Gryseels, Herwig Leirs

## Abstract

In Africa, indigenous multimammate mice (*Mastomys natalensis*) only appear to live commensally in houses when invasive black rats (*Rattus rattus*) are absent, yet little is known about the underlying mechanism. Avoidance through smell may cause the absence of *M. natalensis* from areas occupied by *R. rattus*, but this hypothesis has not yet been tested. We conducted a Y-maze choice experiment where 15 *M. natalensis* were offered a choice between corridors containing conspecific scent, *R. rattus* scent, and a control scent. Residence time in the *R. rattus* corridor was greater than that in the control corridor but equal to that in the *M. natalensis* corridor, suggesting that multimammate mice do not actively avoid the scent of their invasive competitor.

## Introduction

The black rat or roof rat (*Rattus rattus* Linnaeus, 1758) is a household and agricultural pest, a feral invader of natural habitats and a reservoir of zoonotic diseases (Amori and Clout 2003; Meerburg et al. 2009; Aplin et al. 2011; Morand et al. 2015). In sub-Saharan Africa the black rat encounters the smaller, indigenous multimammate mouse (*Mastomys natalensis* Smith, 1834), another important agricultural pest (Makundi et al. 1999; Mulungu et al. 2014) and carrier of zoonotic diseases (Isaäcson 1975; Wulff et al. 1977; Meerburg et al. 2009). *M. natalensis* is a generalist species occurring in most habitats throughout sub-Saharan Africa except dense forests and deserts (Leirs 2013). While it lives commensally in houses in most parts of its western range (Misonne 1959; Duplantier and Granjon 1988; Leirs 2013), it is rarely encountered inside houses in coastal West Africa and throughout eastern Africa, even though it can occur at high population sizes peridomestically and in agricultural fields (Leirs 2013).

This spatial variation in commensalism appears to be linked to the distribution of *R. rattus* (Veenstra 1958; Misonne 1959; Monadjem et al. 2011), which in Africa is almost exclusively commensal. *R. rattus* invaded western Africa only in the 15^th^ century (Rosevear 1969). It is still largely restricted to the coastal regions and the invasion of inland regions is ongoing since a few decades only (Fichet-Calvet et al. 2005; Diagne et al. 2016). In eastern Africa the coastal invasion took place several centuries earlier through the Indo-Pacific trade (Tollenaere et al. 2010). The inland invasion already started in the 19^th^ century (Dieterlen 1979) and *R. rattus* is now widespread from the east coast up to the Democratic Republic of Congo, Zambia and South Africa (Happold 2013). Misonne (1959) describes the invasion of *R. rattus* in villages on the left bank of Lake Albert during the 1950’s, where *M. natalensis* used to be the sole commensal rodent until it was completely displaced in the villages that were first invaded. Notably however they did co-occur in villages that were invaded at a later time (Misonne 1959) and in another study area at Lake Edward, although that study was unclear about whether they co-occurred in the same houses (Misonne 1959).

While it is therefore likely that *R. rattus* can indeed exclude *M. natalensis* from the commensal habitat, it is not clear through which mechanism(s). Several studies suggested that the larger and possibly more aggressive black rat might prevent the multimammate mouse from entering houses through interference competition over space and resources (Veenstra 1958; Misonne 1959; Monadjem et al. 2011). The black rat aggressively defends its territory against conspecifics (Ewer 1971) and shows aggression towards heterospecifics (Stokes et al. 2012; Bridgman et al. 2013). Because such an encounter is potentially harmful to the smaller and less aggressive multimammate mouse (Borremans et al. 2014; Veenstra 1958), the development of avoidance behaviour would not be unexpected. As territorial boundaries and resource ownership in rodent communities are generally mediated by olfactory communication (Heavener et al. 2014), scent would be an ideal indirect signal to allow avoidance. Indeed, some rodents show avoidance in response to scents from heterospecific competitors (Heske and Repp 1986; Krasnov and Khokhlova 1996; Simeonovska-Nikolova 2007).

We conducted a standard Y-maze choice experiment to investigate whether the multimammate mouse actively avoids the black rat on the basis of scent. We hypothesized that multimammate mice would spend significantly less time in a black rat scent corridor than in corridors with a neutral or conspecific scent.

## Methods

### Experimental animals

In July 2014, 15 multimammate mice (10 males and 5 females) were trapped in a maize field on the campus site of Sokoine University of Agriculture, Morogoro, Tanzania, using live traps (LFA; Sherman Live Trap, Tallahassee, FL) baited with peanut butter and maize flour. The mice were individually housed to avoid scent information gathering prior to the experiment. Black rats were housed together in order for the bedding to have a strong scent and to avoid individual effects. Standard rodent cages were used, and food and water were provided ad libitum during the entire experiment. The trapping period coincided with the *M. natalensis* breeding season (Leirs et al. 1994). Avoidance behaviour towards competing rodents might be strongest during this period, as observed for several other rodent species (Heske and Repp 1986; Simeonovska-Nikolova 2007).

### Experimental setup

The experiment was conducted in an acrylic glass Y-maze consisting of three arms (30 cm x 10 cm x 60 cm; height x width x length) connected by a central area that could be closed off from the arms by acrylic glass partitions. The setup was covered with a thick red foil to disturb the animals as little as possible during the observation; as mice have low sensitivity to red light, they perceived this environment as a dark space (Lyubarsky et al. 1999).

Each arm contained one of three sources of wood shavings that were used as bedding: (1) from two cages with conspecifics (one male and one female, randomly chosen, to avoid sex and individual effects); (2) from a cage with two black rats, trapped in a hay barn at the same campus (unfortunately data on sex, weight or sexual maturity status of the rats were lost); (3) unused bedding material as a control. In each experimental run, bedding was placed at the end of a randomly chosen corridor. A multimammate mouse was placed in the centre of the Y-maze, at this point closed off from the corridors. The mouse was left in this closed-off centre for five minutes for acclimatisation to the new environment. The corridors were then opened simultaneously to allow free movement for a period of five minutes, during which time spent in each corridor was measured. After each run, the setup was cleaned thoroughly with soap and rinsed with water to remove all scents. In this way 15 mice were each tested thrice over a period of four days.

### Statistical analysis

The proportion of time spent in each corridor was used as a proxy for preference/avoidance. To determine whether there was a significant difference in residence time between different treatments, likelihood ratio testing of generalized linear models with a binomial distribution (logit link function) was used, with time (i.e. residence time divided by the sum of the residence times spent in the three corridors) as response variable and treatment as fixed effect. We also tested for effects of individual ID and day (the first, second or third day for an individual, which is not necessarily the same day for every individual) in separate models to decide whether or not they should be included in the final model.

We tested for interactions between scent type and sex or weight. Due to the small sample size, the weight classes were restricted to ‘light’ (weight below sample median) and ‘heavy’ (weight above median).

In order to increase the statistical power for comparing the two rodent-scented corridors, the models were repeated using normalised data, where time spent in each rodent-scented corridor was divided by the time spent in the control corridor. Here we used ANOVA of linear models.

Because of the relatively small sample size of our experiment (due to logistical limitations), we conducted a statistical power analysis to quantify the sensitivity of the experiment. This was done by repeatedly simulating data, where the existing data structure (variation, sample size, experimental design) was retained while the effect size (difference in means between the two treatments) was changed over a relevant range of values, assuming a P-value of 0.05 as a “statistical significance cut-off” (R code available on request).

All analyses were conducted in R 3.0.2 (R Development Core Team 2016) using package lme4 (Bates et al. 2015).

## Results

### Non-normalized data

For the non-normalized residence data there were no significant effects of individual (χ^2^ = 3.3, df = 14, P = 0.998) or day (χ^2^ = 0.23, df = 2, P = 0.889). Corridor type had a significant overall effect on residence time (χ^2^ = 11.5, df = 2, P = 0.003; Figure 1), which was due to a significantly lower residence time in the control corridor (*M. natalensis* vs. control: odds ratio = 0.19 95%CI 0.06-0.54, χ^2^ = 10.2, df = 1, P = 0.001; *R. rattus* vs. control: odds ratio = 0.24 95%CI 0.07-0.67, χ^2^ = 7.45, df = 1, P = 0.006), whereas no significant difference was observed between the rodent-scented corridors (odds ratio = 1.24 95%CI 0.53-2.93, χ^2^ = 0.24, df = 1, P = 0.63). There were no significant interactions between corridor and sex (χ^2^ = 1.53, df = 2, *P* = 0.47) or weight (χ*2* = 1.4, df = 2, *P* = 0.5).

**Figure 1:**
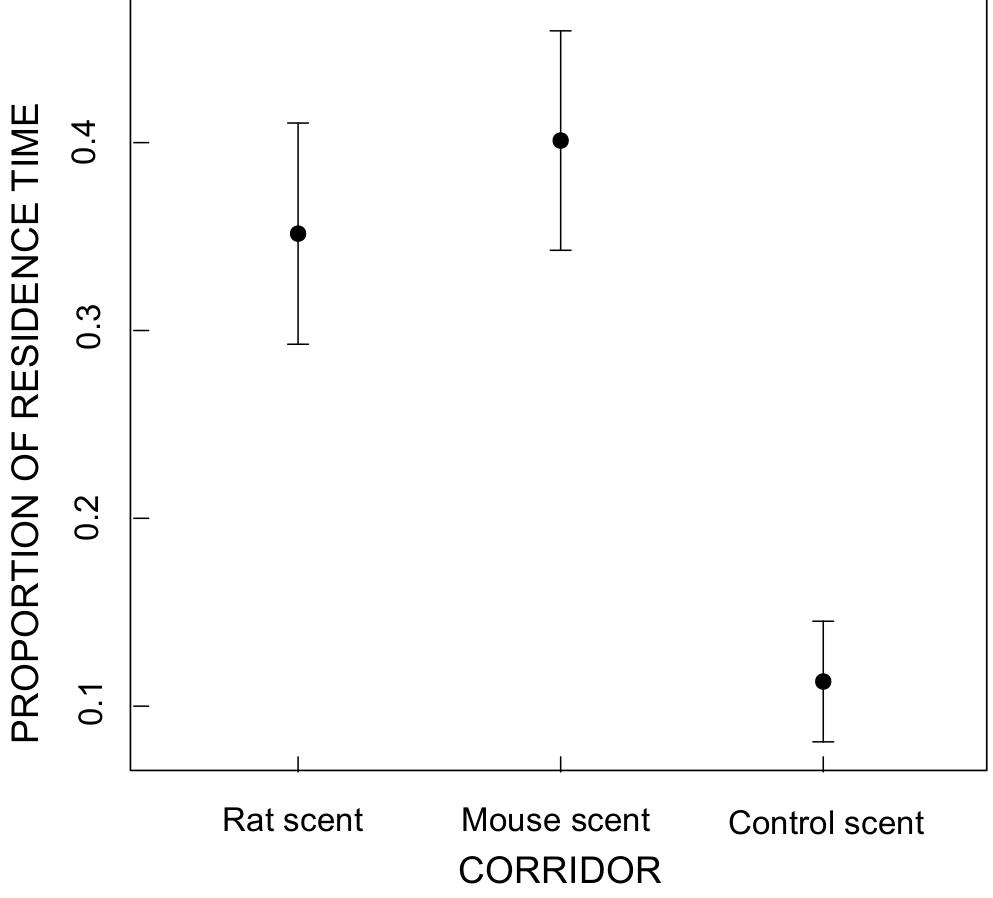
Mean proportion of residence time (± SE) in the three corridors.

### Normalized data

For the normalized data there was a significant effect of day (F = 4.5, df = 2, P = 0.013) but not for individual (F = 1.13, df = 14, P = 0.35), so the former variable was included as a parameter in the final model. There was no significant difference between the two experimental corridors (F = 0.19, df = 1, *P* = 0.6632; Figure 2). There were no significant interactions between corridor and sex (F = 0.02, df = 1, *P* = 0.89) or weight (F = 1.28, df = 1, *P* = 0.26).

**Figure 2:**
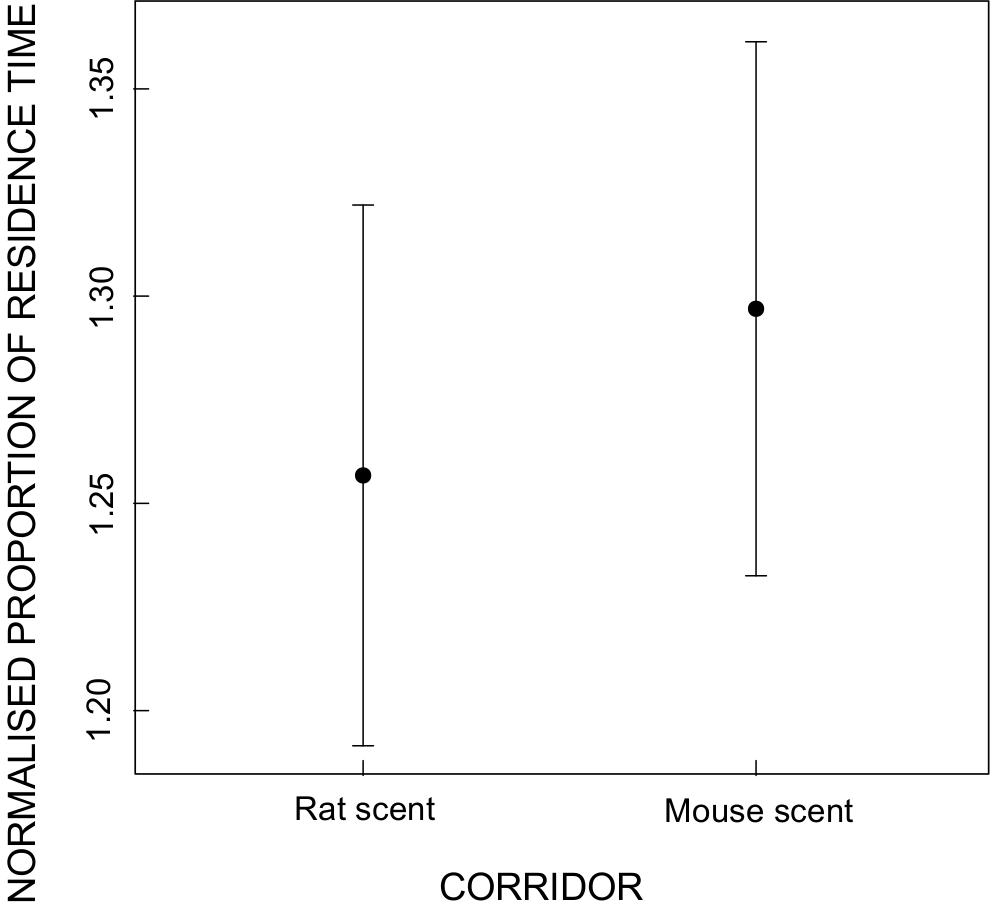
Mean normalised proportion of residence time (± SE) in the two experimental corridors.

The power analysis showed that, given the current sample size and experimental design, it would have been possible to detect a 10% difference between time spent in the two rodent-scented corridors with a probability of 80%, and differences above 12% with a probability of 90% or higher (Figure 3). A difference of 6% would have been detected, but with a probability of 40%. For example, mean residence times of 180 s in one corridor and 202 s in the second would have resulted in a statistically significant difference with a high (90%) probability, while there would be a 40% probability of detecting a statistical difference between 180 s and 191 s.

**Figure 3:**
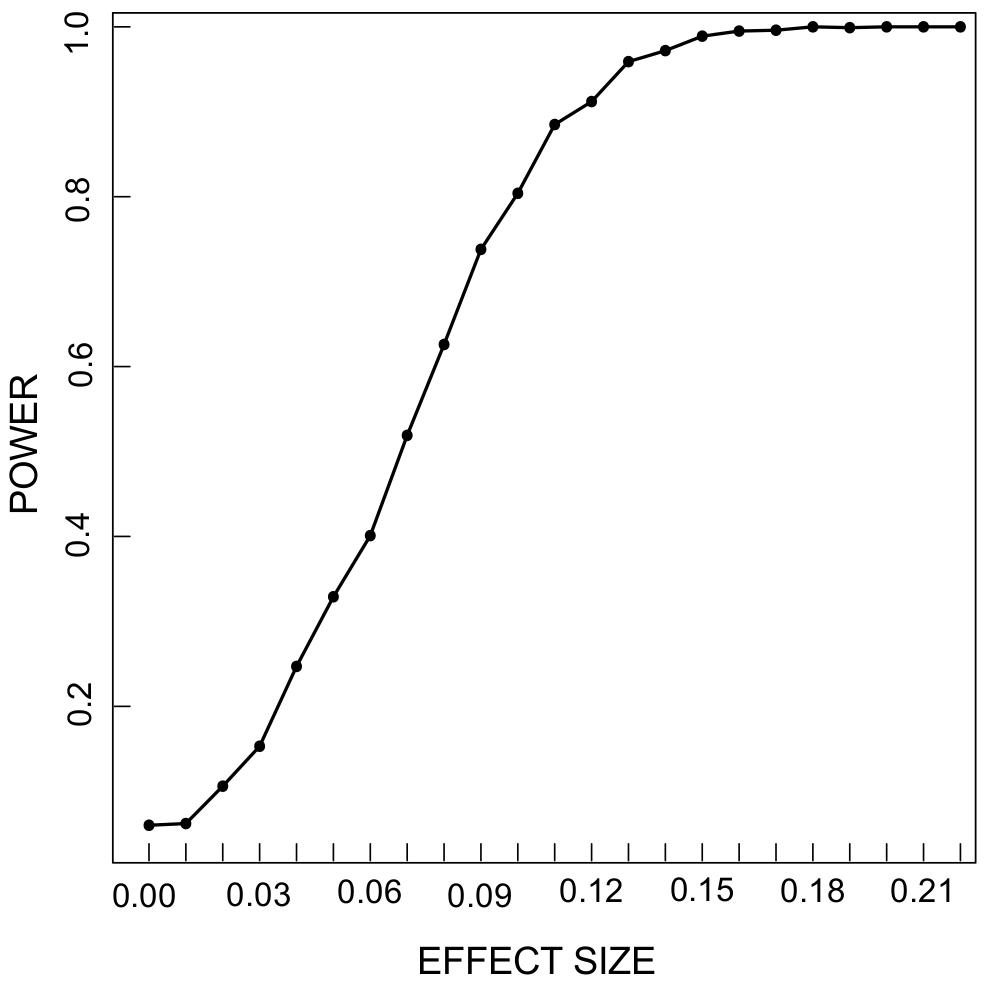
Statistical power for a range of effect sizes.

## Discussion

We tested whether indigenous *M. natalensis* would avoid *R. rattus* based on scent, but found no evidence for a strong effect of this behaviour, as the tested mice spent similar amounts of time in conspecific and black rat corridors. This suggests that multimammate mice do not actively avoid the scent of the invasive black rat. Similar situations have been observed for Cypriot mice (*Mus cypriacus*) and for deer mice (*Peromyscus maniculatus*) that were expected, but not observed, to avoid the scent of competing black rats and to avoid competing and predatory grasshopper mice (*Onychomys leucogaster*) respectively (Stapp and Van Horne 1996; Frynta et al. 2015).

The mice in our study were attracted much stronger to both conspecific scent and rat scent than to the scent of unused bedding material. This was also observed by Frynta et al. (2015), where Cypriot mice and non-commensal Syrian house mice (*Mus musculus*) preferred the scent of other murids over unscented sawdust. While this could be due to explorative behaviour, and interest in the presence of other rodents, another reason for this could be the confounding presence of additional smells (e.g. food) that were present in the bedding of both the *M. natalensis* and *R. rattus,* but not in the control corridor. Nevertheless, if avoidance can be completely overcome by attraction to food in well-fed animals, it is unlikely to be of major importance in real world situations. Avoidance of *R. rattus* by *M. natalensis* through smell would either need to evolve in the *M. natalensis* genetic population, or be learned by individuals through association of the smell with previous negative encounters. The former might indeed not be likely in our study area, as (1) *R. rattus* - *M. natalensis* coexistence history is relatively short (the black rat probably arrived in Morogoro between 260 and 120 years ago (Dieterlen 1979), and (2) because the individual number of *M. natalensis* occupying habitats where they are likely to encounter *R. rattus* is several orders of magnitude smaller than the number of individuals in habitats without *R. rattus*. The selection pressure for evolving *R. rattus* avoidance is therefore small. The second mechanism, i.e. plastically acquired scent avoidance, would require previous encounters with *R. rattus*, and we do not know whether this was the case for the individuals used in our experiments, as they were captured in an agricultural field, where black rat densities are very low (Christensen 1996; Goüy de Bellocq et al. 2010). Testing this hypothesis would require additional experiments on *M. natalensis* that live in close proximity to *R. rattus* (in natural or experimental conditions).

The absence of multimammate mice from habitats occupied by black rats may also be caused by direct competitive interactions that do not involve smell. Although predation by invasive black rats generally does not appear to be a major threat to native mammals (Harris 2009; Smith and Banks 2014), the black rat has been observed to prey on Anacapa deer mice (*Peromyscus maniculatus anacapae*) in California (Ozer et al. 2011) and house mice have been found in its stomach contents in New Zealand (McQueen and Lawrence 2008). Black rats may also repeatedly outcompete multimammate mice from the commensal habitat, either being more effective in occupying shelter and/or consuming food sources, or aggressively defending their territory.

Considering the considerable negative economic impact and public health relevance of both species, there is need for further research into the exact interaction between *M. natalensis* and *R. rattus*.

## Acknowledgments

We are grateful to the Pest Management Centre of the Sokoine University of Agriculture (Morogoro, Tanzania). We thank Shabani Lutea and the Wildlife Management students Abbas Mvungu, Enock Stephano, Salum Mgwama and Zabibu Kabalika for their help during the field and experimental work. This work was supported by the University of Antwerp and the Antwerp Study Centre for Infectious Diseases (ASCID). The authors declare no conflict of interest.

